# Exacerbation of a Subset of Behavioral Phenotypes by Early Treatment with AAV FOXG1 Gene Replacement Therapy in a Mouse Model of FOXG1 Syndrome

**DOI:** 10.1101/2025.10.22.683927

**Authors:** Christina L. Torturo, Kimberly Kerker, Jodi Gresack, Christopher DeGrave, Ailing Du, Dan Wang, Guangping Gao, Scott Reich, Sylvie Ramboz, Dinah W. Y. Sah

## Abstract

FOXG1 syndrome is a severe neurodevelopmental disorder characterized by microcephaly, profound intellectual disability with communication deficits including lack of speech, impaired social interaction, increased anxiety, hyperkinetic/dyskinetic movements, seizures and abnormal sleep patterns. Mutations in a single allele of the *FOXG1* gene cause disease, likely due to loss-of-function. However, current therapies do not target this root cause of FOXG1 syndrome and have little to modest therapeutic benefit on only a small subset of symptoms. Recently, we reported the beneficial effects of adeno-associated virus (AAV) human *FOXG1* gene replacement therapy administered by intracerebroventricular (ICV) injection at postnatal day 6 (P6) on several behavioral deficits that are relevant to key features of human FOXG1 syndrome, in male FOXG1 mice that were engineered with a highly prevalent, patient-specific Q84P mutation. Here, we report the behavioral effects of AAV human *FOXG1* gene replacement therapy administered by ICV injection in female as well as male mice and at an earlier age - postnatal day 2 (P2). Although the earlier studies had suggested that AAV *FOXG1* gene replacement therapy is a promising approach for the treatment of a subset of functional deficits in human FOXG1 syndrome with no toxicity observations, our current study shows that certain motor behaviors can be negatively impacted or exacerbated by P2 treatment with AAV *FOXG1* gene replacement therapy, in female but not male FOXG1 mice. Given these results, the risk-benefit balance of AAV *FOXG1* gene replacement therapy in patients with FOXG1 syndrome should be carefully considered, especially in female patients.

## INTRODUCTION

FOXG1 syndrome is an autism spectrum disorder caused by mutations in a single allele of the *FOXG1* gene. The syndrome is characterized by severe neurodevelopmental deficits that have a profound deleterious impact on the patient’s quality of life. Clinical manifestations include severe global developmental delay and postnatal growth deficiency, resulting in intellectual disability with communication deficits, impaired social interaction, increased anxiety, hyperkinetic/dyskinetic movements, seizures, abnormal sleep patterns, and gastrointestinal and feeding problems (Wong *et al*. 2019). Most symptoms typically begin in the first year of life with diagnoses between birth and 3 years of age (Brimble *et al*. 2023). Current therapies have little to modest therapeutic benefit on only a small subset of symptoms, with no effect on the underlying disease progression. Thus, there is a high unmet medical need for effective treatments that target the root cause of disease.

FOXG1 is a forkhead family transcription factor that is critical for forebrain development and territorial specification, including neurogenesis and balance of excitatory versus inhibitory neuronal differentiation, neurite outgrowth, organization of the cortical layers, and cortico-cortical connections (Wong *et al*. 2019). In addition, FOXG1 plays a key role in dendritogenesis and neural plasticity (Wong *et al*. 2019). Mutations in the *FOXG1* gene that cause FOXG1 syndrome include frameshift, missense, nonsense and deletion variants that result in loss or gain of function (Frisari *et al*. 2022, Brimble *et al*. 2023). A decrease in levels of functional FOXG1 protein disrupts neurodevelopment, leading to abnormalities that include microcephaly, reduced cortical layers and numbers of cortical neurons, atrophy of the hippocampus, agenesis of the corpus callosum and delay in myelination (Pringsheim *et al*. 2019). These structural changes likely contribute to the clinical manifestations.

Human FOXG1 syndrome is believed to be caused primarily by haploinsufficiency. In a mouse model of FOXG1 haploinsufficiency, deletion of one *Foxg1* allele in cortical, hippocampal and hypothalamic neurons and in intermediate progenitor cells (*Foxg1fl/+;NexCre* or *Foxg1*-cHet; Jeon *et al*. 2024) resulted in microcephaly, corpus callosum malformation, reduction of the cortical upper layers, and hippocampus dentate gyrus malformation, similar to structural changes in the human disease. In this mouse model, ICV administration of adeno-associated virus 9 (AAV9) with a human FOXG1 payload driven by a chicken beta-actin (CBA) promoter (AAV9.CBA.hFOXG1) on postnatal day 1 was reported to result in human FOXG1 expression throughout the brain, not only in cells that normally express FOXG1 but also in those that do not normally express FOXG1 (Jeon *et al*. 2024). In conjunction with this expression of FOXG1, some but not all neuroanatomical deficits were ameliorated. The authors reported nearly complete normalization of (A) the increased number of oligodendrocyte progenitor cells (OLIG2+) in the corpus callosum, (B) the reduced number of CTIP2+ cortical neurons (possibly by preventing neuronal cell death), and (C) the shortened length and widened apex angle of dentate gyrus (dentate gyrus abnormalities). Partial normalization of both reduced thickness of the corpus callosum, and reduced myelin basic protein-immunoreactivity in the corpus callosum, reflecting myelination deficits, was also observed. However, there was no effect of AAV9.CBA.hFOXG1 administration on the thickness of the cortex. An important consideration for interpreting results from this *Foxg1fl/+;NexCre* mouse model is that in contrast to human disease, mutant FOXG1 was not present in cortical, hippocampal and hypothalamic neurons or in intermediate progenitor cells, which may impact the phenotype and/or ability to ameliorate the phenotype with AAV9 *FOXG1* gene replacement therapy. Furthermore, it is unclear how the improvement in some but not all neuroanatomical deficits impacts dysfunction, including behavioral deficits, and therefore whether the neuroanatomical effects of AAV human *FOXG1* gene replacement therapy will translate into significant clinical benefit.

Recently, two Q84P mouse models of FOXG1 syndrome have been described (Jeon *et al*. 2025, Torturo *et al*. 2025) that contain a frameshift mutation in one allele of *Foxg1*, resulting in a mutant, truncated form of murine FOXG1 protein. Since a mutant *Foxg1* gene is present, these models more closely represent the genotype of a prevalent form of human FOXG1 syndrome (p.Q86Pfs35X) than the *Foxg1fl/+;NexCre* mouse model which does not contain mutant FOXG1. Both Q84P models recapitulate important functional aspects of FOXG1 syndrome and provide useful models of disease to evaluate effects of potential therapeutics on these behavioral deficits as well as to investigate possible correlations between functional and neuroanatomical changes. In our previous report (Torturo *et al*. 2025), we described the general tolerability and beneficial effects of AAV *FOXG1* gene replacement therapy (scAAV9.hSyn1-opthFOXG1) administered at postnatal day 6 (P6) in male C57BL/6J-Foxg1^em1Mri^ mice (C57BL/6J-Foxg1^Q84P^, courtesy of Dr. Teresa Gunn, McLaughlin Research Institute) on several behaviors related to anxiety and habituation to a novel environment. Here, we describe studies in this Q84P mouse model to evaluate the safety and behavioral efficacy of the same AAV human *FOXG1* gene replacement therapy tested previously, administered again by ICV injection, but at an earlier age – postnatal day 2 (P2), and in female as well as male mice. We focused on P2 for AAV administration to assess possible further benefits of earlier post-natal treatment.

Vector genome (VG) and human *FOXG1* mRNA were quantified to confirm AAV delivery and expression of exogenous human FOXG1 in the brain. To characterize the behavioral phenotype in this mouse model, we evaluated ultrasonic vocalization at 8 weeks of age, and wire hang, open field, tapered balance beam, running wheel, and SmartCube^®^ endpoints at 10 to 13 weeks of age. Compared to vehicle-treated wild-type (WT) mice, there were significant deficits in vehicle-treated Q84P heterozygous mice in ultrasonic vocalization, open field, running wheel and a subset of SmartCube^®^ endpoints, but no differences in wire hang or tapered balance beam endpoints. ICV administration of scAAV9.hSyn1-opthFOXG1 at P2 in Q84P heterozygous mice resulted in significant improvements in ultrasonic vocalization in female mice only, open field center distance traveled in female mice only, running wheel distance traveled on day 1, and multiple SmartCube^®^ deficits. There were no treatment effects on tapered balance beam - latency to turn. However, scAAV9.hSyn1-opthFOXG1 treatment had a detrimental effect on motor function specifically in female but not male mice, as indicated by (A) the wire hang endpoint, reducing the latency to fall, and (B) the tapered balance beam slip-step ratio, increasing the fraction of slips. In addition, scAAV9.hSyn1-opthFOXG1-treated female Q84P heterozygous mice exhibited hyperactivity, indicated by total distance traveled and total rearing frequency in the open field test beyond those in vehicle-treated WT mice. There were no notable effects in Q84P heterozygous mice of scAAV9.hSyn1-opthFOXG1 treatment at P2 on body weight or cage side observations.

Taken together, our results in the Q84P mouse model of FOXG1 syndrome demonstrate that ICV administration of AAV *FOXG1* gene replacement therapy comprising scAAV9.hSyn1-opthFOXG1, with consequent increased levels of human FOXG1 in critical brain regions, ameliorates several but not all behavioral deficits that are relevant to key features of human FOXG1 syndrome. However, importantly, we found that scAAV9.hSyn1-opthFOXG1 administration at P2 had detrimental effects in female mice specifically, including motor dysfunction and hyperactivity. These studies indicate that the risk-benefit balance of AAV *FOXG1* gene replacement therapy in patients with FOXG1 syndrome should be carefully considered, especially in female patients.

## RESULTS

To evaluate the tolerability and behavioral efficacy of AAV *FOXG1* gene replacement therapy, we used the Q84P mouse model of FOXG1 syndrome, heterozygous on a C57BL/6 background. These mice express a murine FOXG1 protein that is both mutated and truncated to 113 amino acids in length, compared to the WT murine FOXG1 protein which is 481 amino acids in length.

### ICV injection of scAAV9.hSyn1-opthFOXG1 in the Q84P mouse on postnatal day 2 did not impact body weight gain or survival

There were no significant differences at any time point during the 14-week study between the body weights of Q84P heterozygous mice treated at P2 with a single ICV dose of vehicle versus those treated with 1.6 × 10^11^ vg scAAV9.hSyn1-opthFOXG1 (**Fig. 1**). However, Q84P heterozygous animals, whether injected with scAAV9.hSyn1.opthFOXG1 or vehicle, weighed significantly less than WT animals injected with vehicle at 4, 5, 6, and 7 weeks of age (**Fig. 1**). In addition, Q84P heterozygous animals treated with scAAV9.hSyn1.opthFOXG1 weighed significantly less than WT animals injected with vehicle at 3, 8 and 9 weeks of age.

**Figure 1.**
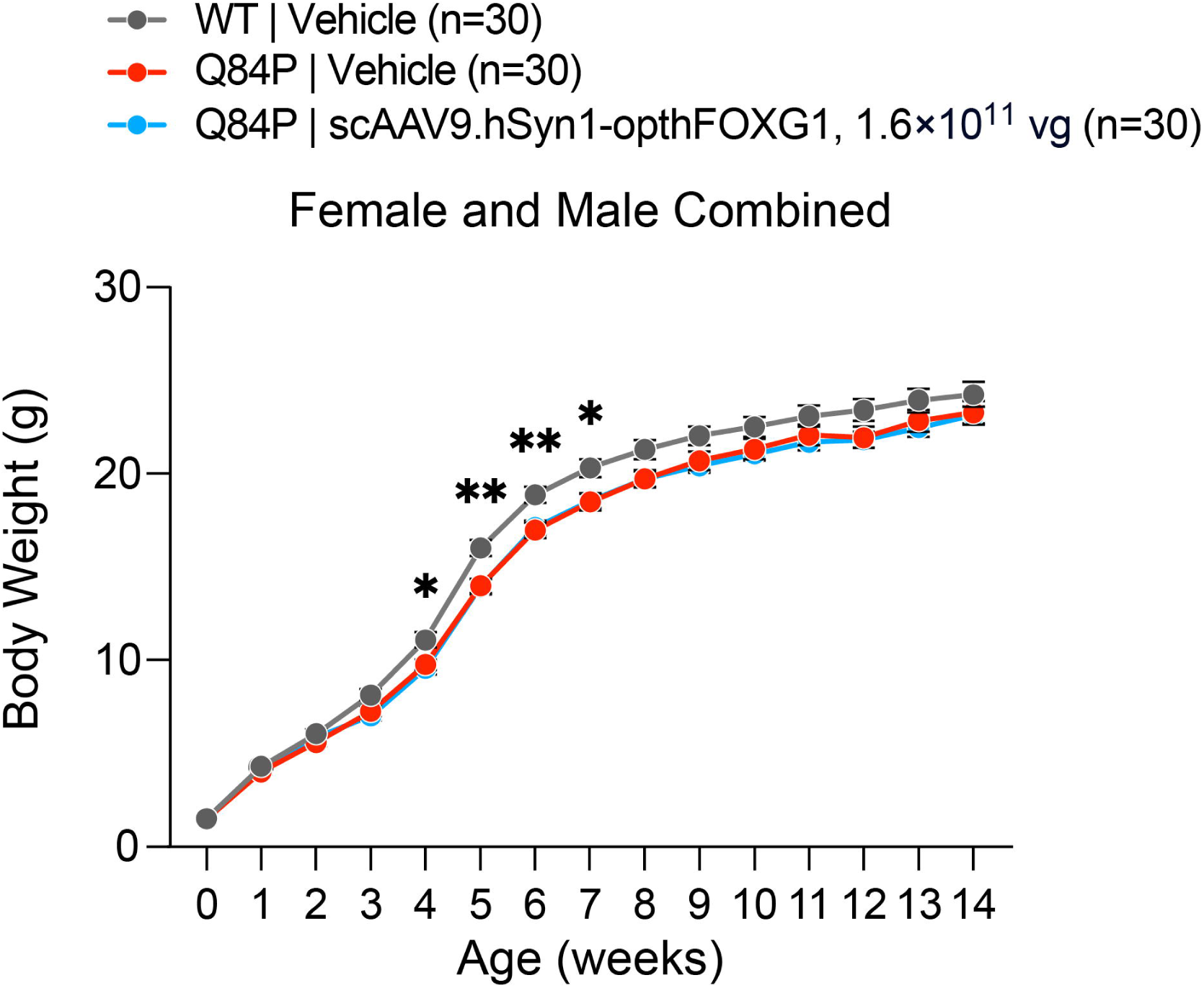
Body weights of Q84P mice treated ICV with scAAV9.hSyn1-opthFOXG1 are not significantly different from Q84P mice treated with vehicle. However, there were significant differences between WT animals treated with vehicle and Q84P heterozygous animals treated with vehicle at 4, 5, 6 and 7 weeks of age. In addition, Q84P heterozygous animals treated with scAAV9.hSyn1.opthFOXG1 weighed significantly less than WT animals treated with vehicle at 3, 4, 5, 6, 7, 8 and 9 weeks of age The number of animals per group (n) represents the number at the time of enrolment. Means ± standard error of the mean are displayed. * p < 0.05, ** p < 0.01, as assessed by two-way ANOVA with Tukey’s multiple comparison test.

All WT and Q84P heterozygous animals survived until the scheduled tissue collection time point at 14 weeks of age with the exception of two WT animals treated with vehicle (at post-natal days 9 and 85), two Q84P heterozygous animals treated with vehicle (at post-natal days 9 and 21) and two Q84P heterozygous animals treated with scAAV9.hSyn1.opthFOXG1 (both at post-natal day 27).

### ICV injection of scAAV9.hSyn1-opthFOXG1 in the Q84P mouse on postnatal day 2 increases FOXG1 expression in key brain regions

VG and human *FOXG1* mRNA levels (**Table 1**) confirmed delivery of scAAV9.hSyn1-opthFOXG1 to the cortex and hippocampus of Q84P heterozygous mice, key brain regions for treating FOXG1 syndrome.

**Table 1.**
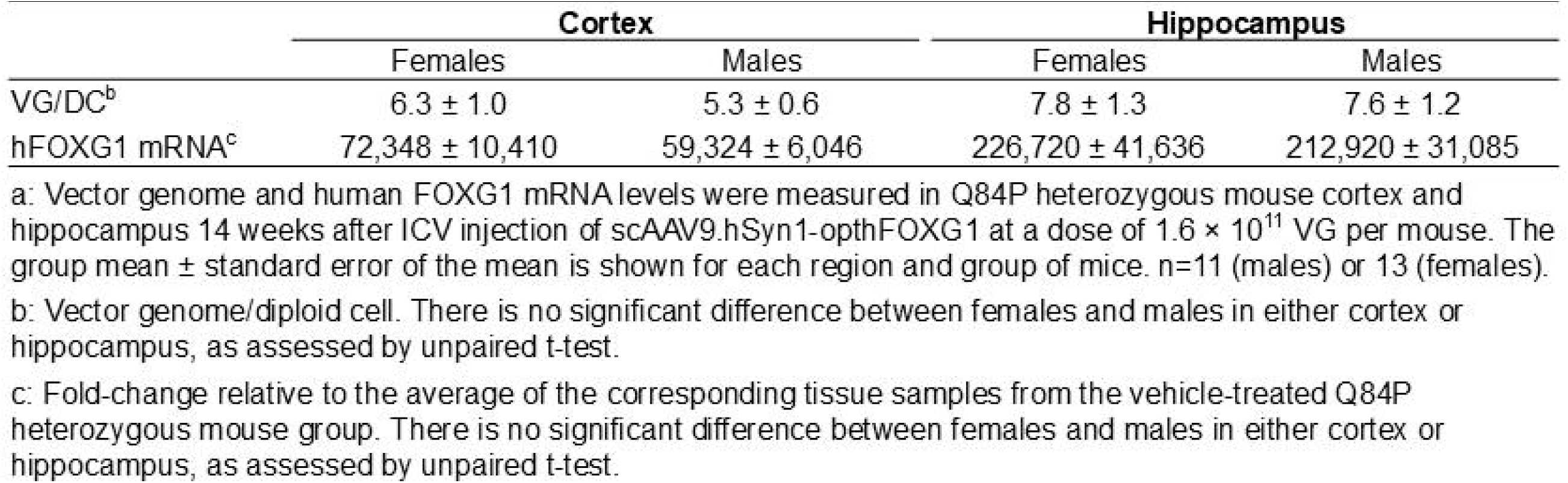
Vector genome and human FOXG1 mRNA levels in Q84P heterozygous mouse cortex and hippocampus•.

VG levels measured by droplet digital PCR (ddPCR) in the Q84P heterozygous mouse cortex and hippocampus at 14 weeks of age after ICV administration of scAAV9.hSyn1-opthFOXG1 were similar in the cortex and hippocampus. There was no significant difference in VG levels between females and males in the cortex or hippocampus. As expected, samples from vehicle-treated WT and Q84P heterozygous animals contained less than 0.01 VG/DC.

Human *FOXG1* mRNA levels in the Q84P heterozygous mouse hippocampus and cortex, quantified by RT-qPCR and normalized to mouse beta-glucuronidase (GUSB) mRNA levels, were also significant after ICV administration of scAAV9.hSyn1-opthFOXG1. Normalized human *FOXG1* mRNA levels, expressed as fold-change relative to the average of the corresponding tissue samples from the vehicle-treated Q84P heterozygous group, were approximately 3-fold higher in the hippocampus than in the cortex. There was no significant difference in normalized human *FOXG1* mRNA levels between females and males in the cortex or hippocampus. In vehicle-treated WT and Q84P mice, human *FOXG1* mRNA levels in cortex and hippocampus, normalized to mouse GUSB mRNA levels, were very low or undetectable as expected since these animals were not administered scAAV9.hSyn1-opthFOXG1.

### ICV treatment with AAV *FOXG1* gene replacement therapy on postnatal day 2 has mixed effects on different behavioral endpoints

#### Ultrasonic vocalization (USV) deficits in Q84P heterozygous mice are improved by AAV FOXG1 gene replacement therapy in females but not males

Overall, the number of USVs at week 8 was lower in vehicle-treated Q84P heterozygous mice compared with WT mice, in both females and males (**Fig. 2A-H**). However, ICV administration of scAAV9.hSyn1-opthFOXG1 at P2 resulted in an increase in the average number of USVs in female mice only, that was significant for continuous calls (**Fig. 2G**) and also observed (but not statistically significant) for total (**Fig. 2A**), short (**Fig. 2C**), complex (**Fig. 2E**) and discontinuous (**Fig. 2G**) calls. In male Q84P heterozygous mice, there were no changes after scAAV9.hSyn1-opthFOXG1 treatment in the number of total (**Fig. 2B**), short (**Fig. 2D**), complex (**Fig. 2F**), or continuous or discontinuous calls (**Fig. 2H**). The duration of USVs was not affected by treatment with scAAV9.hSyn1-opthFOXG1 in either female (**Fig. 2I**) or male (**Fig. 2J**) mice.

**Figure 2.**
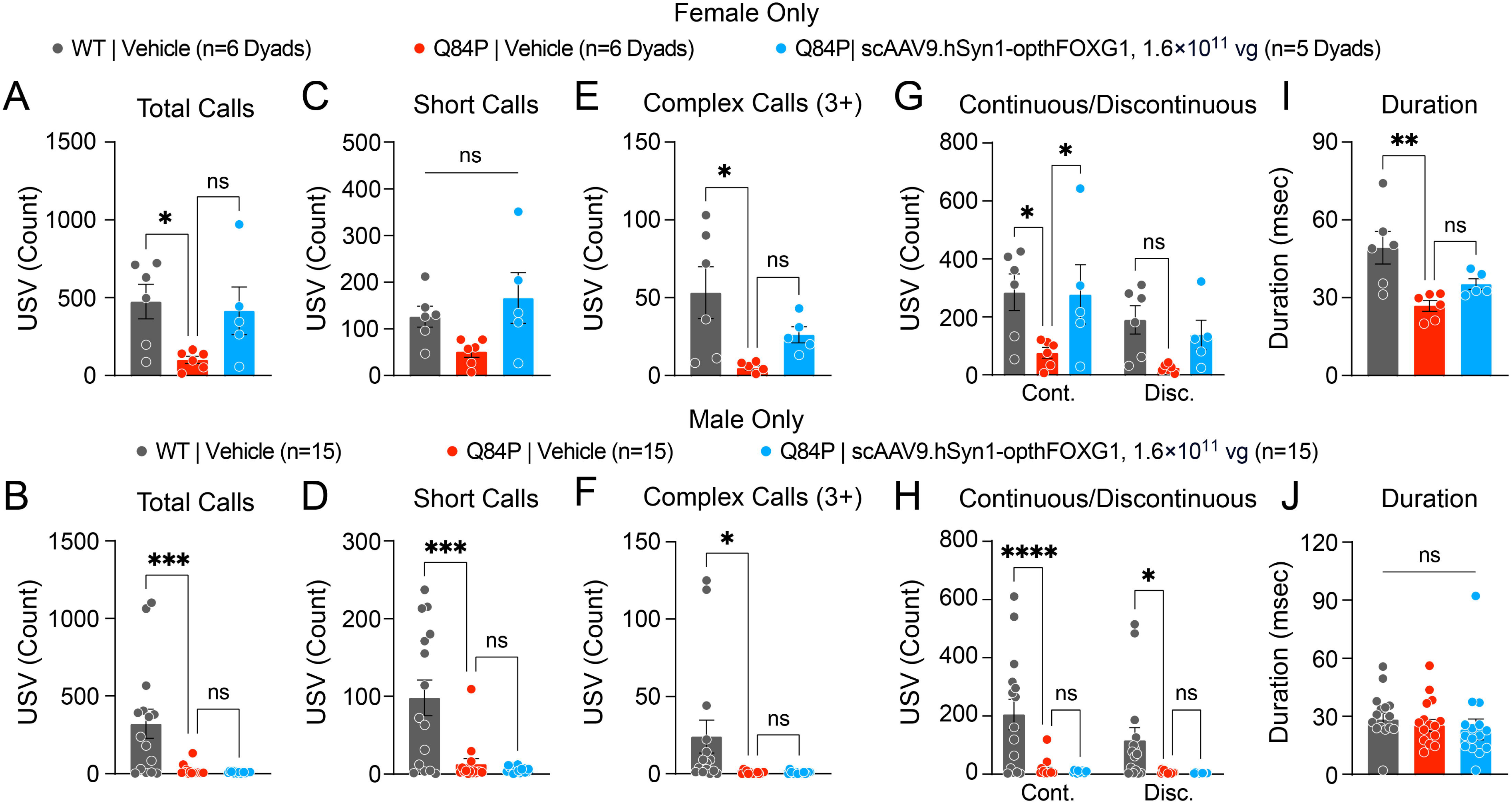
Ultrasonic vocalization (USV) in 8-week old vehicle-treated WT mice, vehicle-treated Q84P heterozygous mice and scAAV9.hSyn1-opthFOXG1-treated Q84P heterozygous mice. Total calls in female (**A**) and male (**B**) mice were significantly different across the treatment groups. Short calls in female (**C**) mice were not significantly different across the treatment groups but were significantly different across the treatment groups in male mice (**D**). Complex (3+) calls in female (**E**) and male (**F**) mice were significantly different across the treatment groups. The number of continuous and discontinuous calls were significantly different across treatment groups in females (**G**) and males (**H**). The duration of USVs was significantly different across the treatment groups in females (**I**) but not in males (**J**). Means ± SEM are displayed. * p < 0.05, ** p < 0.01, *** p < 0.001, **** p < 0.0001, ns: not significant, as assessed by one-way ANOVA with Tukey’s multiple comparison test.

#### Wire hang performance is not changed in vehicle-treated Q84P heterozygous vs WT mice but is worsened by AAV FOXG1 gene replacement therapy in females but not males

There was no difference in wire hang performance between vehicle-treated Q84P heterozygous mice and WT mice (**Fig. 3**). However, after ICV administration of scAAV9.hSyn1-opthFOXG1 at P2, Q84P heterozygous mice performed more poorly with a significantly reduced latency to fall, compared with vehicle-treated Q84P heterozygous or WT mice (**Fig. 3A**). In female Q84P heterozygous mice (**Fig. 3B**), this deleterious effect was significant and more pronounced that in male mice (**Fig. 3C**). These results suggest that AAV *FOXG1* gene replacement therapy administered at P2 induces motor dysfunction, especially in females.

**Figure 3.**
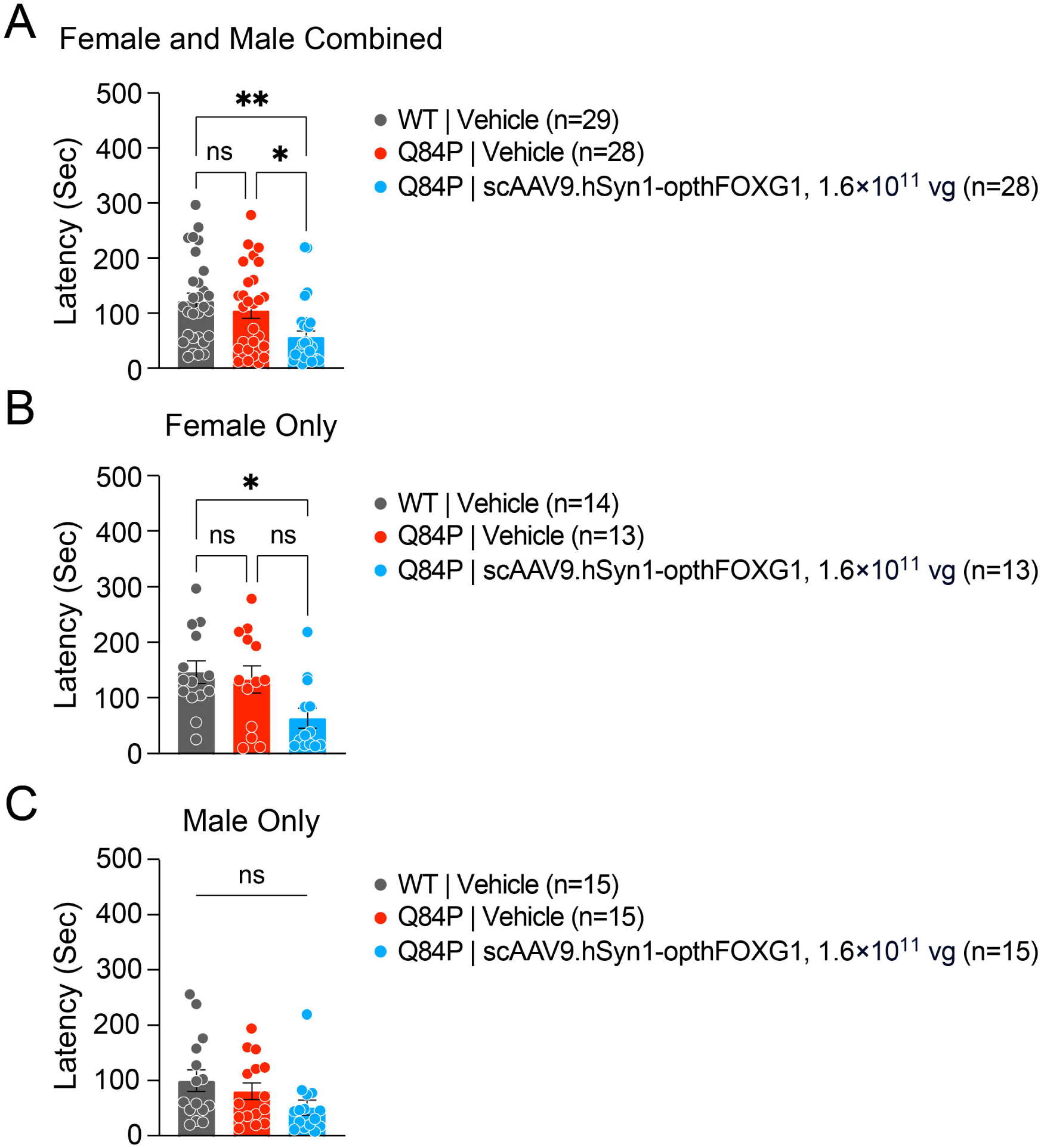
Wire hang latency to fall in 10-week old vehicle-treated WT mice, vehicle-treated Q84P heterozygous mice and scAAV9.hSyn1-opthFOXG1-treated Q84P heterozygous mice. The latency to fall in Q84P mice treated ICV with scAAV9.hSyn1-opthFOXG1 was significantly reduced compared with vehicle-treated WT mice in females but not in males. There were significant differences in latency to fall across the treatment groups with females and males combined (**A**) and in females only (**B**), but not in males (**C**). Means ± SEM are displayed. * p < 0.05, ** p < 0.01, ns: not significant, as assessed by one-way ANOVA with Tukey’s multiple comparison test.

#### Foot slips in the tapered balance beam test were increased by AAV FOXG1 gene replacement therapy in females only

There was no significant tapered balance beam phenotype, comparing latency to turn (**Fig. S1**), latency to traverse (**Fig. S1**) and the ratio of foot slips to steps (**Fig. 4**) between vehicle-treated Q84P heterozygous mice and WT mice. For latency to turn, there was no effect of treatment of Q84P heterozygous mice with scAAV9.hSyn1-opthFOXG1. However, for latency to traverse, scAAV9.hSyn1-opthFOXG1 treatment resulted in a significant increase to a level that was not significantly different from WT mice (**Fig. S1A**).

**Figure 4.**
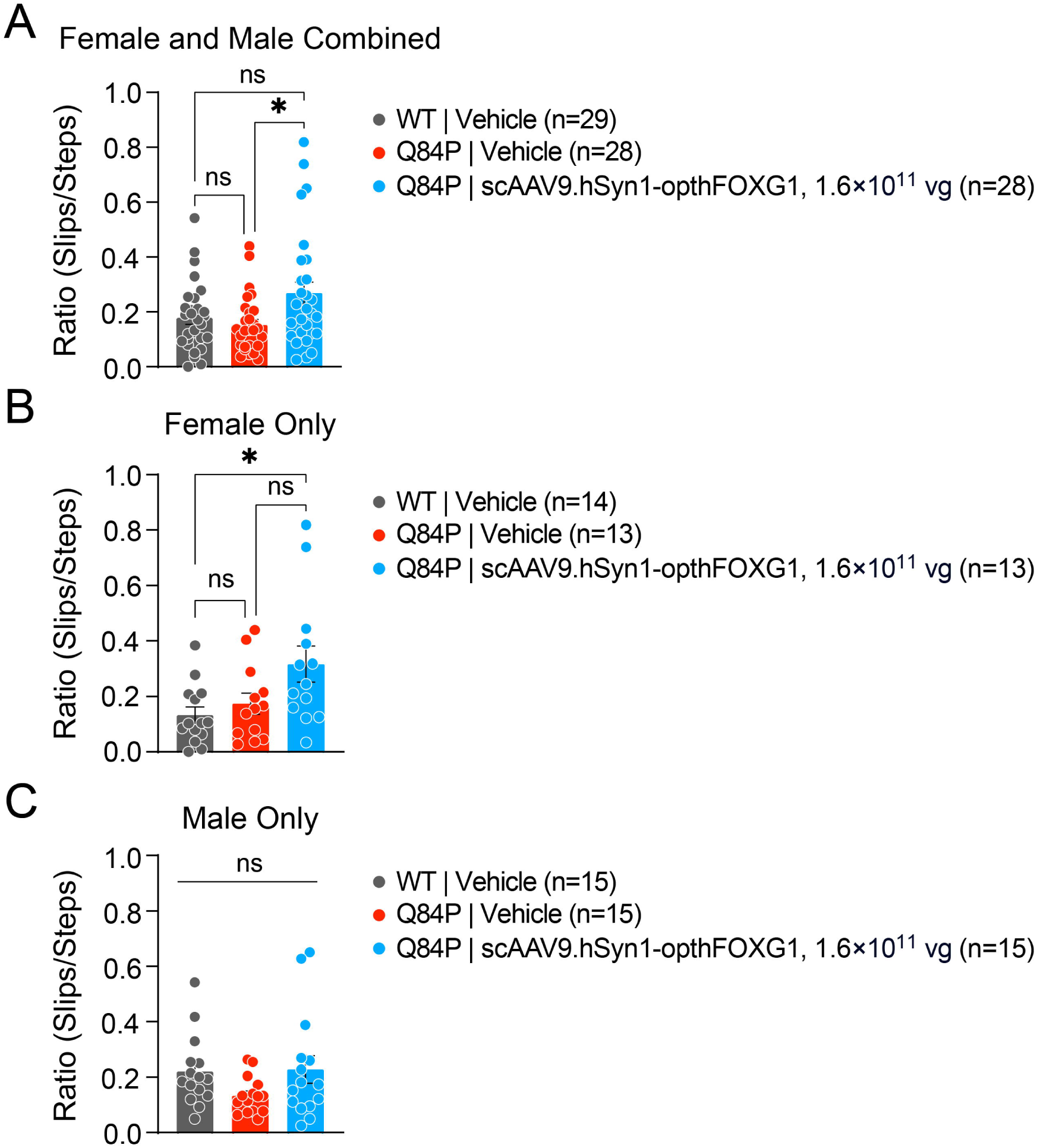
Foot slips in the tapered balance beam assessment in 11-week old vehicle-treated WT mice, vehicle-treated Q84P heterozygous mice and scAAV9.hSyn1-opthFOXG1-treated Q84P heterozygous mice. Foot slips after scAAV9.hSyn1-opthFOXG1 treatment were increased above levels in vehicle-treated WT mice in female Q84P heterozygous mice but not in males. There were significant differences in the ratio of foot slips to steps across the treatment groups with females and males combined (**A**) and in females only (**B**) but not in males only (**C**). Means ± SEM are displayed. * p < 0.05, ns: not significant, as assessed by one-way ANOVA with Tukey’s multiple comparison test.

Notably, after treatment of Q84P heterozygous mice with scAAV9.hSyn1-opthFOXG1, the ratio of foot slips to steps increased significantly compared with vehicle-treated Q84P heterozygous mice (**Fig. 4A**). In scAAV9.hSyn1-opthFOXG1-treated female Q84P heterozygous mice, this ratio of foot slips to steps was significantly greater than in vehicle-treated WT mice (**Fig. 4B**). In contrast, in Q84P heterozygous males, the ratio of foot slips to steps after scAAV9.hSyn1-opthFOXG1 treatment was not significantly different from either vehicle-treated Q84P heterozygous mice or WT mice, indicating that there was no induction of motor abnormality (**Fig. 4C**). These results indicate that AAV *FOXG1* gene replacement therapy administered on P2 induces motor dysfunction in female Q84P heterozygous mice only.

#### Deficits in open field distances traveled were normalized or supra-normalized by AAV FOXG1 gene replacement therapy in females only

Open field total and center distances traveled were lower in vehicle-treated Q84P heterozygous mice vs WT mice (**Fig. 5A** and **D**), which were significant in female Q84P heterozygous mice (**Fig. 5B** and **E**) but not in male Q84P heterozygous mice (**Fig. 5C** and **F**). The total distance traveled was increased significantly after treatment of Q84P heterozygous mice with scAAV9.hSyn1-opthFOXG1, in both females (**Fig. 5B**) and males (**Fig. 5C**). However, in female mice, the total distance traveled after scAAV9.hSyn1-opthFOXG1 treatment was significantly greater than in vehicle-treated WT mice (**Fig. 5B**), indicating a supra-normalization of this behavior. The center distance traveled was also increased after treatment of Q84P heterozygous mice with scAAV9.hSyn1-opthFOXG1, resulting in no significant difference compared with vehicle-treated WT mice in females and males combined (**Fig. 5D**) and in females only (**Fig. 5E**). These results indicate that AAV *FOXG1* gene replacement therapy administered on P2 normalized center distance traveled in female Q84P heterozygous mice, suggesting an amelioration of anxiety, but also induced hyperactivity in total distance traveled in females only.

**Figure 5.**
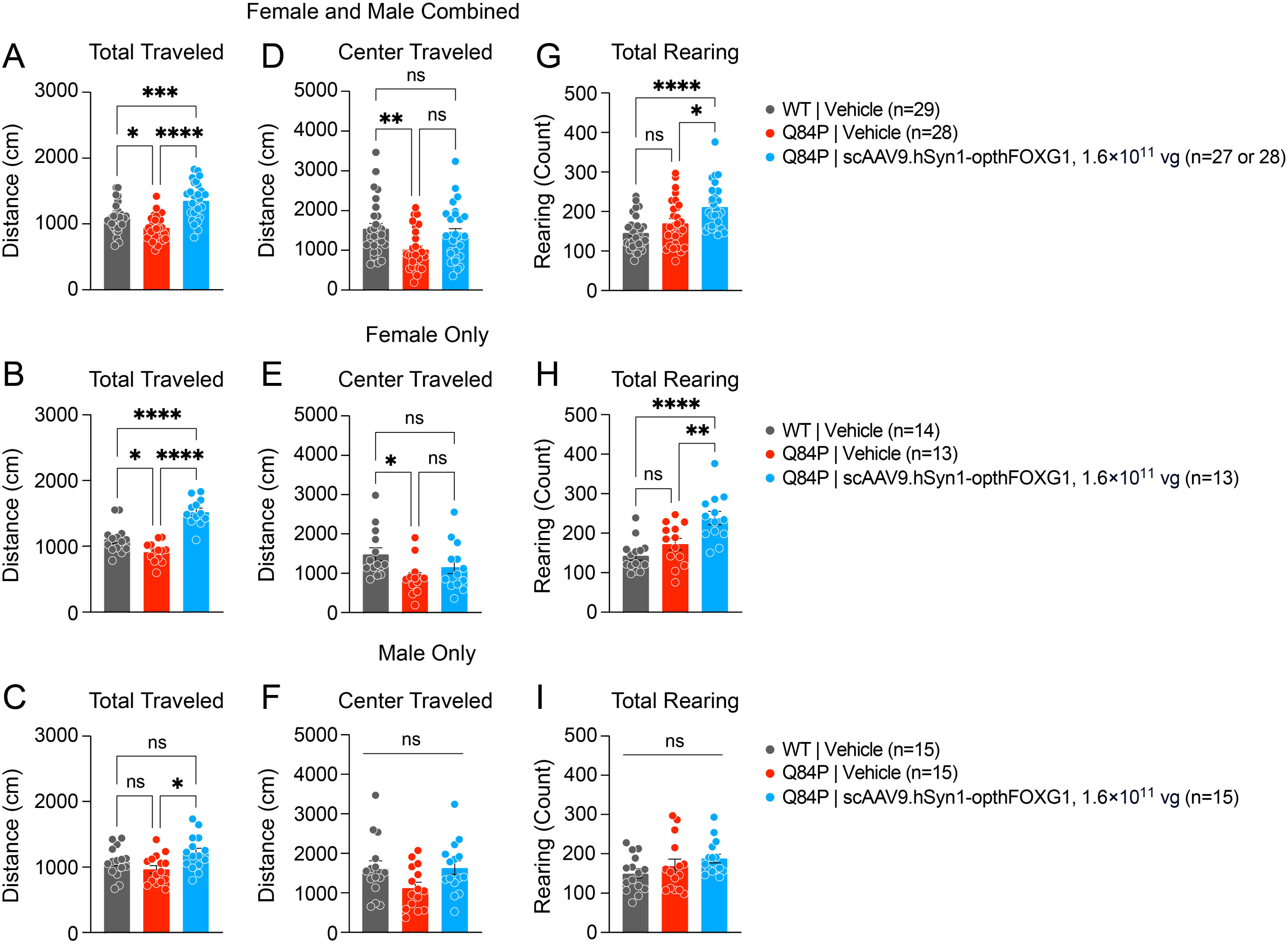
Open field total and center distances traveled and total rearing in 11-week old vehicle-treated WT mice, vehicle-treated Q84P heterozygous mice and scAAV9.hSyn1-opthFOXG1-treated Q84P heterozygous mice. Total and center distances traveled in Q84P mice treated ICV with scAAV9.hSyn1-opthFOXG1 were increased above or to levels, respectively, in vehicle-treated WT mice in females but not in males. There were significant differences in total distance traveled across the treatment groups with females and males combined (**A**), in females only (**B**) and in males only (**C**). For center distance traveled and total rearing bouts, there were significant differences across treatment groups with females and males combined (**D, G**) and in females only (**E, H**) but not in males only (**F, I**). Means ± SEM are displayed. * p < 0.05, ** p < 0.01, *** p < 0.001, **** p < 0.0001, ns: not significant, as assessed by one-way ANOVA with Tukey’s multiple comparison test. For Q84P mice treated ICV with scAAV9.hSyn1-opthFOXG1, female and male mice combined, the number of mice (n) evaluated for total distance traveled was 27 whereas for center distance traveled and rearing bouts, 28 mice were evaluated.

#### Rearing frequency in open field was abnormally increased by AAV FOXG1 gene replacement therapy in females only

There was no difference in open field rearing frequency between vehicle-treated Q84P heterozygous mice and WT mice (**Fig. 5G**). However, after treatment of Q84P heterozygous mice with scAAV9.hSyn1-opthFOXG1, the total rearing count increased in females (**Fig. 5H**) but not males (**Fig. 5I**), and to a level that was significantly greater than in vehicle-treated WT mice. These results indicate that AAV *FOXG1* gene replacement therapy administered on P2 induced hyperactivity in females only.

#### Running wheel activity deficient on Day 1 of assessment is partially normalized by AAV FOXG1 gene replacement therapy

Running wheel activity showed significant reductions during the animals’ light cycle (from 7AM-7PM) and dark cycle (from 7PM-7AM) in both female and male Q84P mice treated with vehicle versus WT mice treated with vehicle (**Fig. 6**). These reductions were present during the light cycle on the first, second and third days of assessment but not on the fourth day of assessment (**Fig. 6A**). The reduction in running wheel activity on the first day of assessment was a robust effect and also significant when females and males were evaluated separately (**Fig. 6B** and **C**, respectively). However, the difference observed on the second day of assessment was statistically significant in males but not females, and on the third day of assessment not statistically significant in either females or males. There was a significant restoration of running wheel activity on day 1 of assessment after treatment of Q84P mice with scAAV9.hSyn1-opthFOXG1 compared with vehicle (**Fig. 6A**, females and males combined). However, there was no significant effect of scAAV9.hSyn1-opthFOXG1 treatment on the day 2 or 3 running wheel deficit. Notably, day 1 is a time period in the running wheel test that is associated with acclimating to and exploring a novel environment (Santos *et al*. 2022). These results suggest that AAV *FOXG1* gene replacement therapy administered on P2 improves habituation to and exploring a new environment, similar to the effect with P6 administration described previously (Torturo *et al*. 2025).

**Figure 6.**
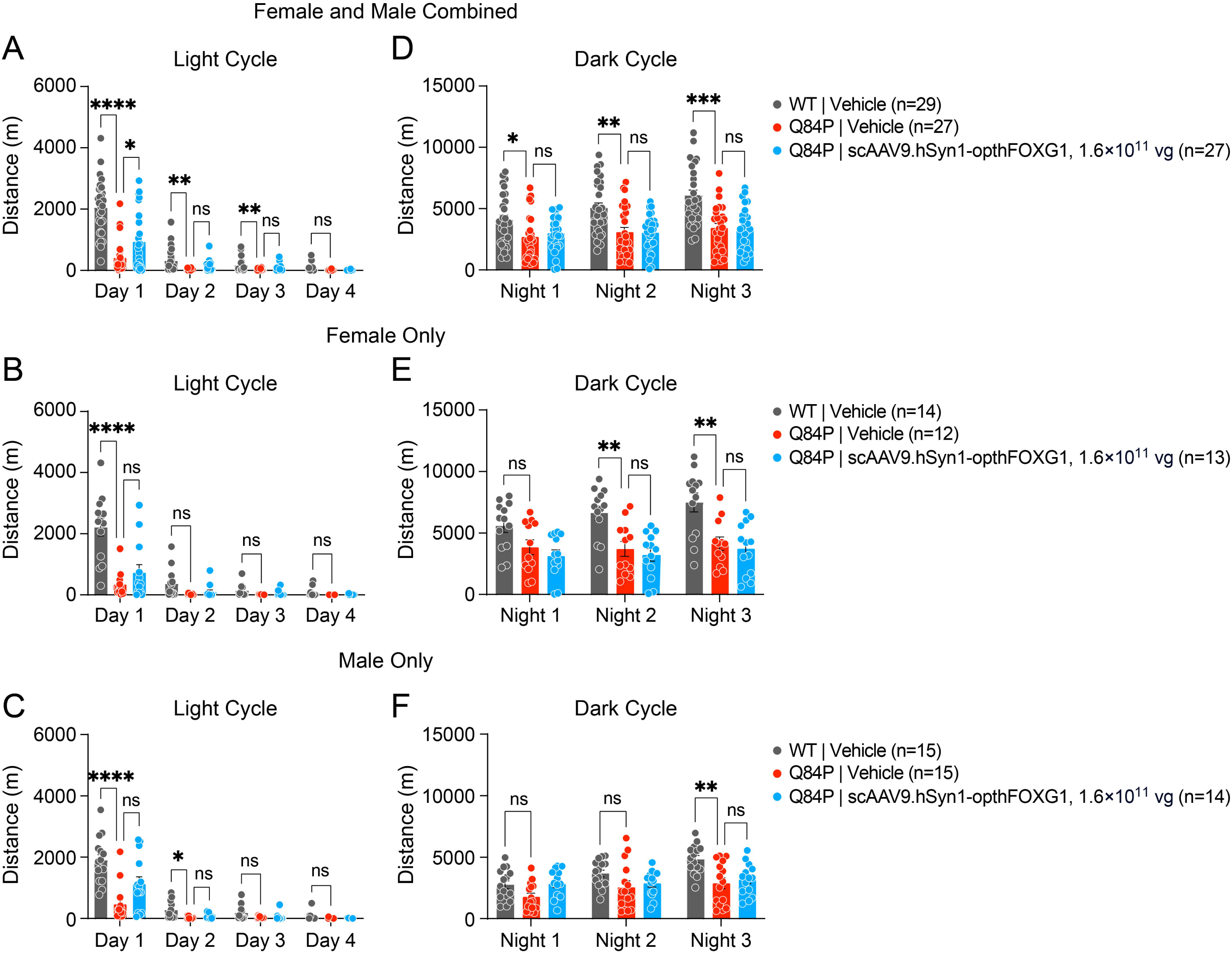
Running wheel total distance traveled during the light (from 7AM to 7PM) and dark (from 7PM to 7AM) cycles in 12-week old vehicle-treated WT mice, vehicle-treated Q84P heterozygous mice and scAAV9.hSyn1-opthFOXG1-treated Q84P heterozygous mice. The total distances traveled during the animals’ light and dark cycles, assessed on 4 days and 3 nights, respectively, showed significant differences between the treatment groups with females and males combined (**A, D**), in females only (**B, E**) and in males only (**C, F**). Means ± SEM are displayed. * p < 0.05, ** p < 0.01, *** p < 0.001, **** p < 0.0001, ns: not significant, as assessed by two-way ANOVA with Tukey’s multiple comparison test.

During the dark cycle, Q84P mice treated with vehicle also traveled significantly less than WT mice treated with vehicle on the first, second and third nights of testing (**Fig. 6D**). However, when females and males were evaluated separately (**Fig. 6E** and **F**, respectively), the difference observed on the first night of assessment was no longer statistically significant and the difference on night 2 was significant in females but not males. The difference on night 3 remained significant in both females and males evaluated separately. There was no significant effect of scAAV9.hSyn1-opthFOXG1 treatment on running wheel distance on any of the nights tested.

In summary, the Q84P mice traveled significantly less on the running wheel than WT mice treated with vehicle on days 1, 2 and 3 of assessment and nights 1, 2 and 3 of assessment. There was a beneficial effect of AAV *FOXG1* gene replacement therapy with scAAV9.hSyn1-opthFOXG1 in Q84P heterozygous mice on day 1 of assessment only, with a partial normalization of running wheel distance traveled.

#### Several SmartCube® behavioral endpoints are improved by AAV FOXG1 gene replacement therapy whereas others are supra-normalized

Phenotypic profiling in SmartCube® provides an assessment of approximately 2,000 features per animal per timepoint, where the features include spontaneous behaviors and responses to challenges. Over the course of the 45-minute testing session, a total of more than 100,000 features are evaluated. At 13 weeks of age, there were significant differences between Q84P heterozygous and WT mice treated with vehicle (**Fig. 7A**). Overall, significant recovery was detected in Q84P heterozygous mice treated with scAAV9.hSyn1-opthFOXG1 compared with Q84P heterozygous mice treated with vehicle (**Fig. 7A**).

**Figure 7.**
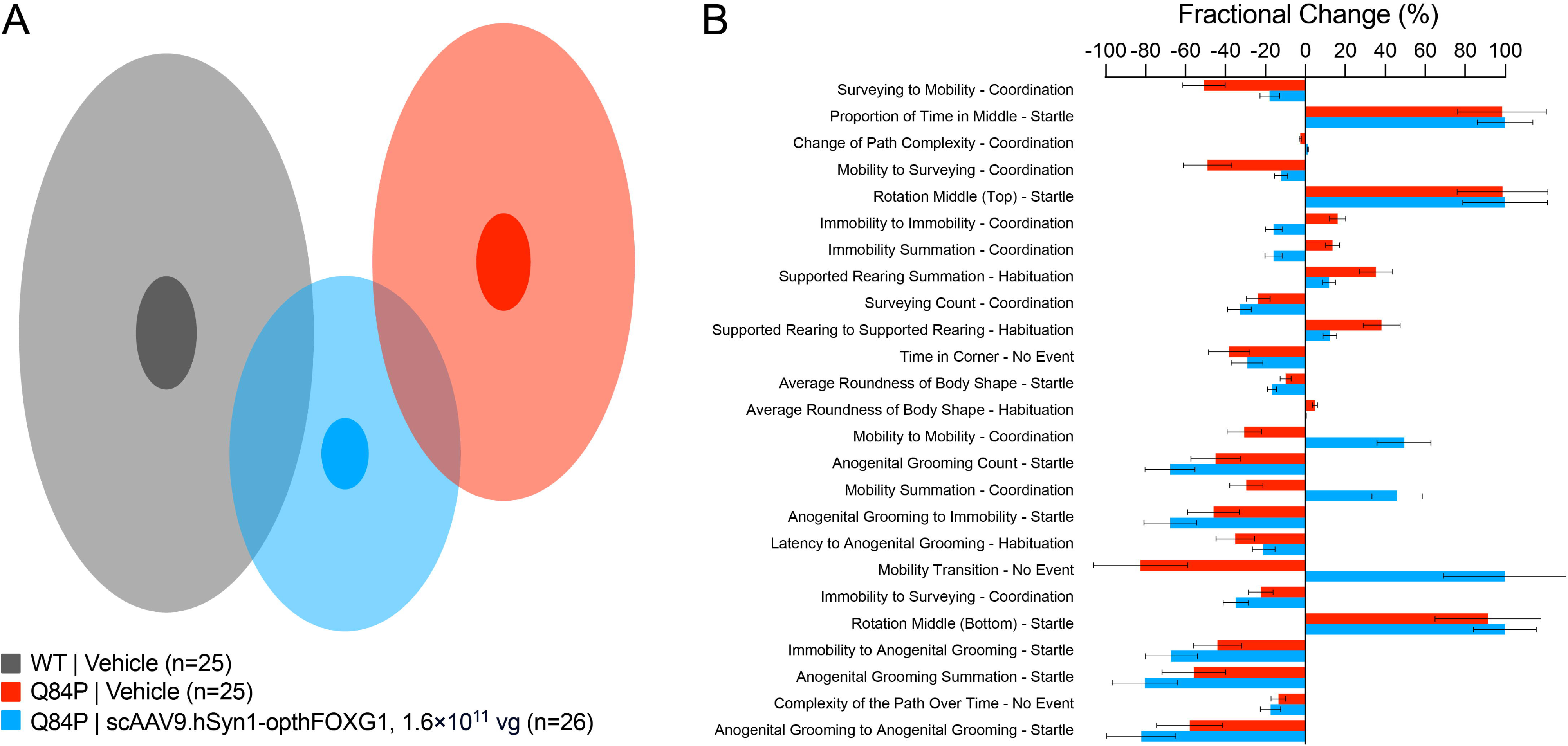
SmartCube^®^ spontaneous behaviors and responses to challenges in 13-week old vehicle-treated WT mice, vehicle-treated Q84P heterozygous mice and scAAV9.hSyn1-opthFOXG1-treated Q84P heterozygous mice. (**A**) DRFA (de-correlated ranked feature analysis) cloud analysis. There were significant differences between Q84P and WT mice treated with vehicle (Discrimination = 83.5%, p < 0.0001) and significant recovery in Q84P mice treated with scAAV9.hSyn1-opthFOXG1 versus vehicle (Recovery = 47.0%, p = 0.0018). (**B**) Twenty-five top features from the SmartCube^®^ results shown as fractional (%) change versus the vehicle-treated WT group for the vehicle-treated Q84P mice (red bars) and for the Q84P mice treated with scAAV9.hSyn1-opthFOXG1 (blue bars). Larger absolute fractional changes indicate greater differences from the vehicle-treated WT group whereas smaller absolute fractional changes indicate smaller differences from the vehicle-treated WT group. SmartCube^®^ data were lost from three vehicle-treated WT mice, three vehicle-treated Q84P heterozygous mice, and two scAAV9.hSyn1-opthFOXG1-treated Q84P heterozygous mice. Means ± standard error of the mean are displayed.

Of the 25 top features of the SmartCube® (**Fig. 7B**) that were significantly different between Q84P heterozygous and WT mice treated with vehicle, four features were substantially improved by treatment with scAAV9.hSyn1-opthFOXG1: Surveying to Mobility – Coordination, Mobility to Surveying – Coordination, Supported Rearing Summation – Habituation, and Supported Rearing to Supported Rearing – Habituation. However, five features were supra-normalized in that a change in one direction (e.g. reduction or increase) comparing scAAV9.hSyn1-opthFOXG1-treated Q84P heterozygous mice versus vehicle-treated WT mice was altered to a change in the other direction (e.g. increase or reduction, respectively): Immobility to Immobility – Coordination, Immobility Summation – Coordination, Mobility to Mobility – Coordination, Mobility Summation – Coordination, and Mobility Transition – No Event.

In general, the features that were improved by AAV *FOXG1* gene replacement therapy with scAAV9.hSyn1-opthFOXG1 reflect an improvement in habituation to a novel environment, whereas those that were supra-normalized reflect coordination.

## DISCUSSION

To assess the therapeutic potential of AAV *FOXG1* gene replacement therapy for treating FOXG1 syndrome, it is critical to establish both safety and functional efficacy in an animal model of disease. Here, in a patient-specific mouse model of FOXG1 syndrome that expresses Q84P mutant FOXG1, we show that AAV *FOXG1* gene replacement therapy to deliver human FOXG1 to neurons in P2 mice had mixed effects. Although AAV *FOXG1* gene replacement therapy ameliorated multiple behavioral deficits that collectively reflect improvements in habituation to a novel environment and reduction of anxiety, deleterious effects such as motor dysfunction and hyperactivity were also observed. Specifically, ICV administration of scAAV9.hSyn1-opthFOXG1 at P2 in Q84P heterozygous mice had beneficial effects on the following behavioral endpoints: ultrasonic vocalization (number of calls in females only), open field (center distance traveled in females), running wheel (only day 1 of assessment) and several SmartCube^®^ assessments. Importantly, although there were no notable effects on body weight or cage side observations, deleterious effect in females only were observed in the following tests: wire hang (decrease in latency to fall), tapered balance beam (increase in footslips) and open field (excess total distance traveled and excess rearing frequency). The tapered balance beam assessment is a particularly sensitive measure of motor impairment and capable of detecting motor deficits early and even when other measures of motor function, such as the rotarod, fail to show any deficits (Heng *et al*. 2007; Brooks *et al*. 2011). Notably, there was a clear sex dependence of the behaviors that were deleteriously affected by ICV administration of scAAV9.hSyn1-opthFOXG1 at P2, which were present only in female Q84P heterozygous mice and not in males.

Vector genome (VG) and human *FOXG1* mRNA levels in cortex and hippocampus, two key brain regions relevant to FOXG1 syndrome, were substantial after ICV administration of 1.6 × 10^11^ vg scAAV9.hSyn1-opthFOXG1 in Q84P heterozygous mice at P2. There were approximately 3-fold higher levels of human *FOXG1* mRNA in the hippocampus than in the cortex whereas VG levels were only slightly higher in the hippocampus than in the cortex. Importantly, there were no sex differences in VG or human *FOXG1* mRNA levels in the cortex or hippocampus after scAAV9.hSyn1-opthFOXG1 treatment. Thus, the deleterious effects of scAAV9.hSyn1-opthFOXG1 treatment in Q84P heterozygous mice specific to females cannot be attributed to different levels of human *FOXG1* expression in the cortex or hippocampus compared with males.

In the previous study (Torturo *et al*. 2025), we evaluated the behavioral effects of ICV administration of 4 × 10^10^, 1 × 10^11^ and 2.6 × 10^11^ vg scAAV9.hSyn1-opthFOXG1 in older Q84P heterozygous mice (P6) and only in males, in contrast to the current study with administration of 1.6 × 10^11^ vg scAAV9.hSyn1-opthFOXG1 in P2 Q84P heterozygous mice, both females and males. A comparison of the behavioral effects in the current versus previous study is informed by the levels of human *FOXG1* expressed. The VG levels in the present study (females and males combined) were approximately 3-fold lower than those in male Q84P heterozygous mice administered 2.6 × 10^11^ vg scAAV9.hSyn1-opthFOXG1 in the earlier study: 5.8 ± 0.6 and 7.7 ± 0.9 VG/DC in the present study versus 17 ± 5.0 and 22 ± 6.3 VG/DC in the previous study, in the cortex and hippocampus, respectively. A similar difference was present when VG levels in only males from the present study were compared with male Q84P heterozygous mice administered 2.6 × 10^11^ vg scAAV9.hSyn1-opthFOXG1 in the earlier study. However, in these same groups of mice, the human *FOXG1* mRNA levels in the present study (females and males combined) were similar to those in the previous study: 66,379 ± 6,304 and 220,395 ± 26,164 -fold (normalized to vehicle-treated Q84P heterozygous group) in the present study versus 47,217 ± 10,486 and 376,511 ± 112,841 -fold in the previous study, in the cortex and hippocampus, respectively. This similarity was maintained when human *FOXG1* mRNA levels in only males from the present study were compared with male Q84P heterozygous mice administered 2.6 × 10^11^ vg scAAV9.hSyn1-opthFOXG1 in the earlier study.

Despite the similar human *FOXG1* mRNA levels in the cortex and hippocampus across the two studies, a comparison of the behavioral effects of P2 treatment in females and males in the present study versus P6 treatment in males in the earlier study revealed pronounced differences. In the open field test of total distance traveled, treatment at P6 with 2.6 × 10^11^ vg scAAV9.hSyn1-opthFOXG1 normalized the reductions observed in vehicle treated Q84P mice relative to WT mice (Torturo *et al*. 2025), whereas treatment at P2 in the present study increased the total distance traveled beyond that in WT mice in females only. In the open field test of total rearing frequency, treatment at P6 with 2.6 × 10^11^ vg scAAV9.hSyn1-opthFOXG1 increased the rearing count significantly to levels that were not different from those in vehicle treated WT mice (Torturo *et al*. 2025). However, treatment at P2 in the present study increased the total rearing frequency to levels that were significantly higher than those in vehicle treated WT mice, in females only. Thus, the excessive total distance traveled and excessive total rearing frequency represented different behavioral effects between the 2 studies that were specific to scAAV9.hSyn1-opthFOXG1 treatment at P2 of female Q84P heterozygous mice; these effects were not observed in male Q84P heterozygous mice treated with scAAV9.hSyn1-opthFOXG1 at P2 or P6.

In an earlier study of the *in vivo* efficacy of AAV gene therapy in a mouse model of FOXG1 haploinsufficiency that does not express mutant FOXG1 (*Foxg1fl/+;NexCre* mouse, Jeon *et al*. 2024), neuroanatomical effects in the brain were reported. ICV administration of AAV9.CBA.hFOXG1 on postnatal day 1 corrected or improved some neuroanatomical deficits in the brain as assessed on postnatal day 27, increasing the reduced numbers of neurons in the cortex and oligodendrocyte progenitor cells in the corpus callosum, increasing the reduced myelination in the corpus callosum, and improving hippocampus dentate gyrus structural abnormalities. However, there was no change in the decreased thickness of the cortex. The improvements in neuroanatomical deficits, while encouraging, do not address the effect of AAV *FOXG1* gene replacement therapy on functional endpoints. Moreover, the lack of mutant FOXG1 in the *Foxg1fl/+;NexCre* mouse may impact the effect of a potential therapeutic approach. For example, if there is a dominant negative effect of mutant FOXG1, then the *Foxg1fl/+;NexCre* mouse would not enable an appropriate evaluation of the effectiveness of a potential therapeutic approach.

The present study with ICV administration of scAAV9.hSyn1-opthFOXG1 at P2 in a patient-specific Q84P mouse model that expresses mutant FOXG1 demonstrated improvements in some behavioral deficits, no change in others, and supranormalization of others. Importantly, motor dysfunction, detected in specific behavioral tests used in the present study but not in the previous study (Torturo *et al*. 2025), emerged in females only after ICV administration of scAAV9.hSyn1-opthFOXG1 at P2 in Q84P heterozygous mice. In particular, the motor deficits were revealed only in female mice and with the wire hang and tapered balance beam assessments, which were not evaluated in the previous dose-response study (the previous single dose study did not contain a sufficient number of animals for assessment of effect). These results underscore the importance of using specific behavioral assessments in a mouse model of disease to evaluate tolerability, rather than relying only on neuroanatomical endpoints. Furthermore, it is unknown whether treatment of female Q84P heterozygous mice with scAAV9.hSyn1-opthFOXG1 at P6 would result in motor deficits or not, since only male Q84P heterozygous mice have been evaluated with treatment at P6 (Torturo *et al*. 2025).

Elevated levels of FOXG1 expression are associated with an autistic spectrum disorder similar to FOXG1 syndrome (Yeung *et al*. 2009, Brunettie-Pierri *et al*. 2011). Therefore, it is particularly important to ensure that there are no detrimental effects of AAV *FOXG1* gene replacement therapy, which may result in long-term levels of exogenous human FOXG1 that are significantly higher than normal. In our studies with 13 weeks of observation in heterozygous Q84P mice following ICV administration of scAAV9.hSyn1-opthFOXG1, the potential risks associated with continuous elevated levels of FOXG1 expression, especially related to motor function, need to be explored further with comprehensive toxicity studies together with quantitation of levels of exogenous FOXG1 protein in specific brain regions for correlation with safety and any adverse findings.

In addition, a dominant-negative effect of mutant Q84P FOXG1 protein would have important implications for therapeutic approaches to treating FOXG1 syndrome, in that elevating levels of WT FOXG1 protein may or may not be sufficient to completely overcome the effects of the mutant protein. It remains to be determined if other behavioral deficits can be addressed by replacement of missing FOXG1 protein earlier in development or in other cell types, or may require suppression of mutant Q84P FOXG1 protein.

In summary, our experiments with ICV administration of scAAV9.hSyn1-opthFOXG1 at P2 in a patient-specific mouse model of FOXG1 syndrome that contains a mutant Q84P *Foxg1* allele in place of a WT *Foxg1* allele, demonstrate the potential therapeutic benefit as well as safety risks of AAV *FOXG1* gene replacement therapy in neurons for the treatment of FOXG1 syndrome. In particular, ICV administration of scAAV9.hSyn1-opthFOXG1 at P2 improved several behaviors related to habituation to a novel environment and anxiety, suggesting that for these behaviors, increasing levels of WT FOXG1 are sufficient to overcome the effects of haploinsufficiency and/or expression of mutant Q84P FOXG1. However, other behaviors were deleteriously affected in female but not male Q84P heterozygous mice, especially those involving motor function. These results suggest that ICV treatment with scAAV9.hSyn1-opthFOXG1 in early post-natal development may have undesirable consequences, especially in females. Further studies in the Q84P mouse model of FOXG1 syndrome to understand the balance of benefit and risk with AAV *FOXG1* gene replacement therapy will be critical for evaluating the translational potential of this therapeutic approach.

## MATERIALS AND METHODS

### Constructs and vectors

scAAV9.hSyn1-opthFOXG1 was produced by the University of Massachusetts Chan Medical School Viral Vector Core. The transgene DNA was packaged in a recombinant AAV9 capsid. Recombinant AAV vectors were produced by triple transfection of adherent HEK293 cells and purified using previously described protocols. In brief, recombinant vectors were purified from clarified lysates using cesium chloride density gradient centrifugation. Vectors were formulated in phosphate-buffered saline (PBS) with 5% sorbitol and 0.001% (w/v) Pluronic F-68^®^, pH 7.2-7.4, and stored below -60°C. Vector titers were measured by ddPCR. For vectors used in vivo, the relative purity of the capsid proteins was confirmed using sodium dodecyl-sulfate polyacrylamide gel electrophoresis (SDS-PAGE) and protein staining. On the day of use, AAV vectors were removed from storage at -60°C, allowed to thaw on refrigerated cold packs, and then maintained at approximately 4°C. Dilutions were made with formulation buffer and then stored at approximately 4°C for no more than 1 week until use. Vehicle groups received formulation buffer.

### Animals and treatments

#### Animals

All animal procedures were approved by the Institutional Animal Care and Use Committee (IACUC).

Q84P heterozygous mice were generated by Dr. Teresa Gunn using CRISPR/Cas9 gene editing (Torturo *et al*. 2025). All mice used in the studies described here were descended from a single male founder and had been backcrossed to wildtype C57BL/6J for at least 5 generations.

Male heterozygous Q84P mice were received at PsychoGenics from Dr. Teresa Gunn and bred with WT C57BL/6 female mice. At birth, a tail snip sample was taken for genotyping to identify heterozygous or WT animals to be placed on study. All animals were examined, manipulated and weighed prior to initiation of the study at postnatal day 6 to assure adequate health and suitability and to minimize non-specific stress associated with manipulation. Mice were maintained on a 12-hour light/dark cycle with food and water available ad libitum.

During the study, animal body weights were recorded daily or twice per week until weaning and twice per week or weekly after weaning, and survival was checked twice per day.

#### Genotyping

Genomic DNA was isolated from neonate tail clippings using Zymo Quick DNA 96 Plus Kit (Cat# D4070) according to the manufacturer’s instructions. PCR reactions were conducted as previously described (Torturo *et al*. 2025).

#### Test articles and dosing

Stock solutions of scAAV9.hSyn1-opthFOXG1 (2.69 × 10^13^ GC/mL) and formulation buffer (vehicle: PBS with 5% sorbitol and 0.001% F-68) were stored at -80°C. On the first day of dose administration, test articles were thawed on wet ice. Once thawed, test articles were stored at 4°C and did not undergo any additional freeze-thaw cycles.

For administration of either vehicle or scAAV9.hSyn1-opthFOXG1, animals were anesthetized using cryoanesthesia. ICV injections were administered bilaterally (3 µL/side for P2 mice) with 10 µl Hamilton Syringes (1701 RN Model, Cat# 7653-01) and custom Hamilton 30 ½ g needles (point style 4, bevel 12°, Cat# 7803-07). Animals were then placed on a warm heating pad immediately following the ICV injection. Once all animals from the litter were dosed and warmed, they were returned to the dam.

### Behavioral assessments

#### Ultrasonic vocalization

The USV assessment was performed at 8 weeks of age.

For adult females, three days prior to USV testing, mice were single housed. On the day of USV testing, the female mice were brought to the USV testing room and allowed to acclimate for a minimum of 1 hour. Females were tested for USV in pairs. One female was placed into the cage of another female that she was originally cage mates with (prior to the 3-day period of single housing). The USV recording began as soon as the former female cage mate was placed into the other cage mate’s cage. The USV recording ended after 300 seconds. During the recording sessions, all USVs were recorded from the female pair.

For adult males, mice were housed in a male-only room for a minimum of 1 week prior to USV testing. Two days before the text, male mice were brought to the USV testing room where they acclimated for a minimum of 1 hour. After acclimation, a sexually mature female mouse was placed into each male cage. Males were paired with females for a minimum of 1 hour after which the females were removed from the cages. At the conclusion of the male-female pairing, males were brought back to their male-only colony room. On the day of USV testing, males were brought to the USV testing room and allowed to acclimate for a minimum of 1 hour. A single cage was placed into each soundproof USV detector. A gauze pad moistened with at least 30 µL of fresh sexually mature female urine was placed into the cage. The USV recording session began as soon as the gauze pad had been placed into the cage. The recording session ended after 300 seconds.

#### Wire hang

The wire hang test of motor function was conducted when the animals were 10 weeks of age by following a modified protocol as described in Santa-Maria *et al*. (2012). Mice were placed on top of a standard wire cage lid. The lid was lightly shaken to cause the animal to tighten its grip and then the lid was then turned upside down. The latency of mice to fall off the wire grid was measured, and average values were computed from three trials (30 seconds apart). Trials were stopped if the mouse remained on the lid for 5 minutes.

#### Open field

The open field (OF) test was used to assess both anxiety-like behavior and motor activity at 11 weeks of age. The OF chambers were plexiglass square chambers (27.3 x 27.3 x 20.3 cm; Med Associates Incs., St Albans, VT) surrounded by infrared photobeam sources (16 x 16 x 16). The enclosure was configured to split the open field into a center and periphery zones and the photocell beams were set to measure activity in both zones. Animals having higher levels of anxiety or lower levels of activity tend to stay in the corners of the enclosure. On the other hand, mice that have higher levels of activity and lower levels of anxiety tend to spend more time in the center of the enclosure. Horizontal activity (distance traveled) and vertical activity (rearing) were recorded from consecutive beam breaks. Animals were placed in the OF chambers for 30 minutes and total ambulatory distance traveled, ambulatory distance traveled in the center, and total rearing were quantified during this period. After testing, animals were placed back into their home cage.

#### Tapered balance beam

The tapered balance beam assessment was performed at 11 weeks of age. The balance beam has a strip of smooth black acrylic 100 cm in length, with a square cross section that tapers from a width of 1.5 cm to 0.5 cm. The beam also has a 0.5 cm safety ledge located 2 cm below the beam with the ledge maintaining a constant width of 0.5 cm as the beam tapers. The angle of the beam is 17° from horizontal running from low to high. The highest point of the beam is approximately 58 cm from the floor. At the opposite side of the balance beam (‘end’ portion) there is a goal box constructed from black acrylic, measuring 10.5 cm^3^ and containing a 3 cm² entrance hole.

On the training day, each animal was required to complete 4 traversals. On testing day (24 hours later), mice received 3 trials of testing with an inter-trial interval of 30 seconds. Mice were placed on the bottom of the beam, facing away from the goal box. The time from placement on the beam to turning to face the goal box was recorded as the latency to turn. The maximum amount of time an animal had to complete the turn was 120 seconds. If the animal did not turn during this time, then it was positioned on the beam facing the goal box for the next phase of the experiment. Once the animal was facing the goal box, the latency to traverse the beam was recorded. The maximum amount of time an animal had to complete the traversal was 120 seconds. During the beam traversal, the number of foot slips was recorded. The slip-step ratio was calculated by dividing the total number of slips by the total number of steps taken per limb.

#### Running wheel

Running wheel activity was assessed at 12 weeks of age. The wheels used were from Lafayette Instrument®, Mouse Activity Wheel with Dual Licometer (model 80822). The polycarbonate chambers measured 13.9”L x 9.25”W x 7.875”H (35.3 x 23.5 x 20 cm) and the aluminum running wheels measured 5.0” ID (12.7 cm) by 2.25” (5.72 cm) width (inside) for a Run Distance of 0.40 meters/revolution. The run surface consisted of 38 rods 0.188” diameter on 0.4298” centers with a 0.2418” gap. The metric equivalent is approximately 4.8 mm diameter on 10.9 mm centers with a 6.14 mm gap. The activity wheels were connected to an interface which electronically recorded the animal’s activity (running interval, average speed, and total distance).

Animals were loaded into the chambers containing the running wheels at approximately 11AM on Day 1 of assessment and subsequently removed from the chambers at the same time on Day 4, 72 hours later. Mice remained in the activity wheel chambers 24 hours/day the three consecutive testing days, after which they were returned to their home cages.

#### SmartCube^®^

The SmartCube^®^ assessment was performed at 13 weeks of age. SmartCube^®^ is a proprietary platform developed by PsychoGenics that employs computer vision to detect changes in body geometry, posture, and behavior (both spontaneous and in response to specific challenges).

Mice were taken in their home cage to the SmartCube^®^ suite of experimental rooms where they remained until they were placed in the apparatus. The standard SmartCube^®^ protocol was run for a single session lasting 45 minutes. After the session, mice were placed back into their home cage and were returned to the colony room.

### Plasma, serum, cerebrospinal fluid and tissue collection

All biological samples were collected upon completion of the in-life phase of the study when the animals were 14 weeks of age.

Animals were anesthetized with isoflurane. CSF was collected by cisternae magna puncture with pulled glass microcapillaries.

After CSF collection, whole blood was harvested via cardiac puncture and processed to plasma using K-EDTA tubes or to serum using serum collector tubes. Plasma and serum samples were frozen on dry ice and then stored at -80°C.

After blood and CSF collection, animals were perfused with ambient temperature PBS to remove blood cells. The brain was then removed and hemisected to collect and snap-freeze the cortices and hippocampi. After snap-freezing, tissue samples were stored at -80°C.

### Vector genome and human FOXG1 mRNA quantitation

#### DNA and RNA isolation

DNA and RNA samples were prepared as previously described (Torturo *et al*. 2025).

#### Vector genome quantitation

Vector DNA levels were quantified using duplex TaqMan-based ddPCR assays. The TaqMan reagents used included mTfrc:TaqMan® Copy Number Reference Assay, mouse, Tfrc (Thermo Scientific), and codon-optimized human FOXG1 (opthFOXG1): forward primer (AGCACAGGCCTGACCTTTAT), reverse primer (CATAGGGTGGCTGGGATAGG), and probe (CCCTGCACCACCCAAGGGCC) (Thermo Scientific). ddPCR was performed using the QX200 Droplet Digital PCR System (Bio-Rad). The number of vector genomes per diploid cell (VG/DC) was calculated by dividing the copies/µL obtained with the opthFOXG1 probe set by that obtained with the mTfrc probe set and then multiplying by two.

#### Human FOXG1 mRNA Quantitation

Human FOXG1 mRNA levels were quantified in cDNA samples using qPCR assays as previously described (Torturo *et al*. 2025).

### Statistical analysis

The unpaired *t* test, or one-way or two-way analysis of variance (ANOVA) was used to assess differences across treatment groups. The Tukey’s multiple comparisons test was employed for post-hoc comparisons after the identification of main effects with ANOVA.

## ACKNOWLEDGMENTS

The authors thank Teresa Gunn for Q84P heterozygous mice; the Viral Vector Core at UMass Chan Medical School for producing the AAV vector; the PsychoGenics team for in vivo assessments; Richard Miller for assistance in quantifying all DNA and RNA samples as well as preparing DNA samples; Meredith McLerie for overseeing early DNA and RNA isolation experimental workflows and designing the experiments for validating the Taqman Primer/probe sets for RT-qPCR multiplexing; and Eric Smith for assistance with the figures.

## SUPPLEMENTAL FIGURE LEGENDS

**Supplemental Figure 1.**
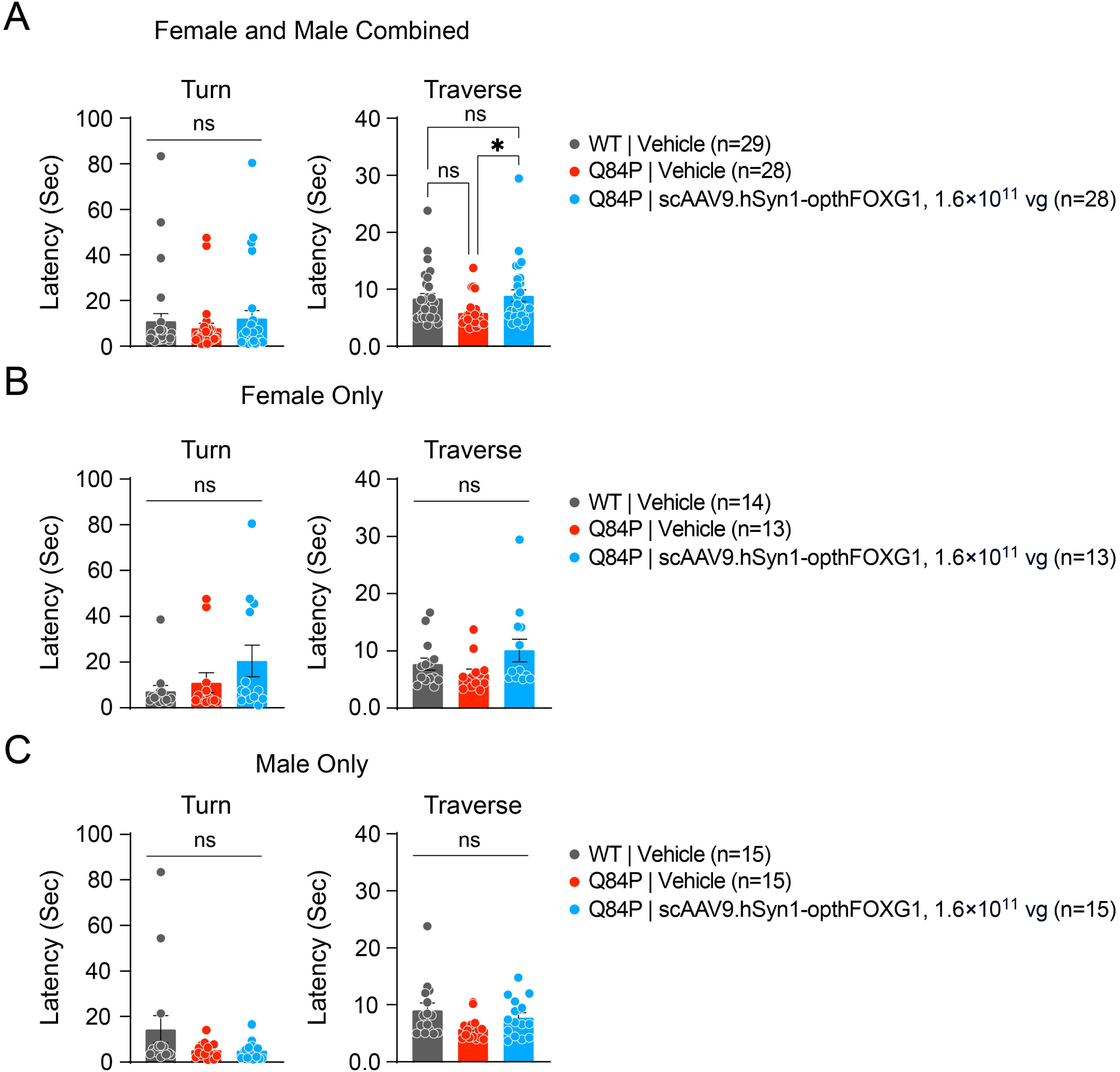
Tapered balance beam latency to turn in 11-week old vehicle-treated WT mice, vehicle-treated Q84P heterozygous mice and scAAV9.hSyn1-opthFOXG1-treated Q84P heterozygous mice showed no significant differences across treatment groups in females and males combined (**A**), females only (**B**) or males only (**C**). Tapered balance beam latency to traverse at 11 weeks of age showed significant differences across treatment groups in females and males combined (**A**) but not in females only (**B**) or males only (**C**). Means ± SEM are displayed. * p < 0.05, ns: not significant, as assessed by one-way ANOVA with Tukey’s multiple comparison test.

